# Patients with peripheral artery disease demonstrate altered expression of soluble and membrane-bound immune checkpoints by peripheral blood immune cells

**DOI:** 10.1101/2024.08.29.610318

**Authors:** R.D. Reitsema, S. Kurt, I. Rangel, H. Hjelmqvist, M. Dreifaldt, A. Sirsjö, A.K. Kumawat

## Abstract

Cardiovascular events seem to occur more frequently after treatment with immune checkpoint inhibitors. This suggests a possible role of immune checkpoints in accelerating atherosclerosis formation. We therefore aimed to assess the expression of soluble and membrane-bound immune checkpoints in patients with peripheral artery disease (PAD).

We assessed the levels of 14 soluble immune checkpoints in fasting blood plasma of PAD patients (intermittent claudication, n= 37) and healthy controls (HCs, n=39) by Multiplex protein assay. We measured the levels of T-cell immunoglobulin and mucin domain 3 (TIM-3), programmed cell death 1 ligand 1 and 2 (PD-L1/2), B-and T-lymphocyte attenuator (BTLA), programmed cell death 1 (PD-1), glucocorticoid-induced TNRF-related protein (GITR), herpesvirus entry mediator (HVEM), cytotoxic T-lymphocyte associated protein 4 (CTLA-4), indoleamine 2,3-dioxygenase (IDO), CD28, CD27, CD80, CD137 and lymphocyte-activation gene 3 (LAG-3). The surface expression of CD28, PD-L2, PD-1, GITR, LAG-3, TIM-3 and BTLA on peripheral blood immune cells was determined by flow cytometry. Cytokine production capacity was measured by flow cytometry in TIM-3+ T cells to determine immune exhaustion.

Soluble levels of PD-L2 were decreased in PAD patients, but only in females, whereas soluble levels of TIM-3 showed a trend towards increased concentration in female PAD patients. All monocyte subsets had increased frequencies of PD-L2+ cells in PAD patients. CD4+ T cells had increased frequencies of TIM-3+ cells, which showed little overlap with other immune exhaustion markers. TIM-3+ CD4+ T cells appeared to have a low capacity to produce pro-inflammatory cytokines, but a higher capacity to produce IL-10 compared to TIM-3-CD4+ T cells.

We found that PAD patients show differences in expression of both membrane-bound and soluble immune checkpoints. Some of these differences might be caused by prolonged immune activation, although immune exhaustion markers did not always overlap.

## Introduction

Peripheral artery disease (PAD) is an atherosclerotic disease that is strongly ageing related. PAD is a significant contributor to cardiovascular morbidity and mortality and is believed to affect more than 230 million people worldwide. PAD affects arteries that perfuse the lower limbs, which leads to claudication and pain, resulting in loss of mobility and quality of life. Loss of lower limbs caused by impaired blood flow is one of the most severe outcomes of PAD (1). PAD is characterized by three different stages, including the asymptomatic stage, the intermittent claudication (IC) stage and the critical limb ischemia (CLI) stage. All stages of PAD, including the asymptomatic stage, are additionally associated with increased cardiovascular disease mortality and morbidity, compared to patients with stable coronary artery disease (CAD) and other forms of cardiovascular disease (CVD) (2).

PAD is caused by atherosclerosis, which is now being recognized as a chronic inflammatory disease affecting the arterial wall (3,4). The first steps of the inflammatory pathway include endothelial dysfunction and the entering and retention of monocytes in the sub-endothelial space. Transformation of monocytes into macrophages is being followed by an immune response against oxidized lipoproteins. Macrophages accumulate lipids intracellularly, which results in the induction of foam cells and more pro-inflammatory cytokine secretion. This leads to the formation of a necrotic core in the vascular wall, which consists of lipids and debris. T cells are involved in the formation of the atherosclerotic plaque as well, as the adaptive immune response against for instance oxidized lipoproteins leads to the aggravation of the pro-inflammatory immune response (5,6).

Immune checkpoints are key regulators of the immune response. Co-inhibitory immune checkpoint receptor-ligand interactions lead to inhibition of the immune response, whereas stimulatory receptor-ligand interactions activate the immune response. The most well-known immune checkpoint interaction involves the expression of CD28 on T cells and their ligands CD80/CD86 on antigen presenting cells (APCs), which is crucial in activating naïve T cells upon antigen encounter (7). Inhibitory checkpoints such as programmed death-1 (PD-1) are often targeted in immunotherapy to treat cancer. By removing this natural brake on the immune system, T cells are being reactivated which is beneficial for anti-tumor immune responses. Not surprisingly, by inhibiting co-inhibitory immune checkpoints, immune-related adverse events can occur related to excessive activation of the T cell response. For instance, cardiovascular events are now being associated with the use of immune checkpoints inhibitors (5,8). This leads to the hypothesis that proper immune regulation by immune checkpoints is crucial in preventing atherosclerotic diseases.

Therefore, we aim to assess the expression of soluble and membrane-bound immune checkpoints in PAD patients. Soluble immune checkpoints are either produced by mRNA expression or are cleaved-of forms of membrane-bound checkpoints and can be measured in plasma. They can interfere with membrane-bound immune checkpoint interactions and either positively or negatively affect these interactions (9). The presence of soluble immune checkpoints therefore forms an additional level of immune regulation. Soluble immune checkpoints can also serve as biomarkers for disease severity. Some checkpoints, such as the co-inhibitory checkpoint and marker for immune exhaustion lymphocyte-activation gene 3 (LAG-3), is associated with clinical characteristics and disease outcome such as in advanced head and neck cancer (10). In cardiovascular disease, an elevation of the stimulatory checkpoint Glucocorticoid-induced TNFR-related protein (GITR) plasma levels was found compared to healthy controls (HCs) (11).

After assessing the levels of soluble immune checkpoints using a commercially available Multiplex panel of 14 immune checkpoints, we further investigated the membrane-bound expression of several of these immune checkpoints. We included GITR based on previous findings in cardiovascular disease that were mentioned above. We also investigated the expression of the exhaustion markers/ co-inhibitory checkpoints T cell immunoglobulin and mucin domain 3 (TIM-3), LAG-3 and B- and T-lymphocyte attenuator (BTLA). TIM-3 was previously found to be an important CD8+ T cell regulator in atherosclerosis (12). The LAG-3 gene is associated with increased risk for myocardial infarction and BTLA pathway stimulation led to reduced lesion development in atherosclerotic mice models (13,14). We included CD28 as a marker of immune maturation and senescence, as lack of CD28 expression is associated with ageing and pro-inflammatory cytokine production. As our study found differences regarding soluble PD-L2 concentrations, we also investigated the PD-1/PD-ligand 2 (PD-L2) axis in T cells and APCs, respectively. In summary, this study will provide a more comprehensive overview of immune checkpoint expression in PAD patients.

## Material and methods

### Study population

The study group was part of the PIMM (PAD inflammation, microbiome and metabolites) study at the School of Medical Sciences, Örebro University, Sweden (Table 1). Symptomatic PAD patients with intermittent claudication, were recruited at the Department of Cardio-thoracic and Vascular Surgery, University Hospital Örebro (USÖ), Sweden, between October 2021 and December 2023. Healthy controls (HCs) matched by sex and age (+/- 5 years) were recruited at the School of Medical Sciences. Participants with dementia, autoimmune and other inflammatory diseases such as kidney disease, diabetes, irritable bowel syndrome, inflammatory bowel disease, and participants having an active infection, having received antibiotics within two months of study enrolment or any over the counter or prescriptive probiotic or bowel cleansing preparation within the past two months of study enrolment were excluded in this study.

**Table 1:** Clinical characteristics of the study population.

|  | Soluble immune checkpoints |  |  | Membrane-bound immune checkpoints |  |  |
| --- | --- | --- | --- | --- | --- | --- |
|  | HC (n=39) | PAD (n=37) | p-value | HC n=27 | PAD n=24 | p-value |
| **Demographics** |  |  |  |  |  |  |
| Age, median (years) | 72.0 | 77.0 | 0.001 | 72 | 76 | 0.057 |
| Sex, M/F | 19/20 | 20/17 |  | 14/13 | 14/10 |  |
| Active smoker, n | 2 | 12 |  | 1 | 7 |  |
| Previous smoker, n (quit > 1 year) | 16 | 17 |  | 11 | 11 |  |
| Non-smoker, n | 21 | 7 |  | 15 | 5 |  |
| **Clinical parameters (median)** |  |  |  |  |  |  |
| ABI right | 1.21* | 0.72 | <0.001 | 1.22* | 0.73 | <0.001 |
| ABI left | 1.20* | 0.66* | <0.001 | 1.21* | 0.66* | <0.001 |
| Intima Media Thickness, Max (mm) | 0.89† | 1.04* | 0.001 | 0.88† | 1.04* | 0.009 |
| Intima Media Thickness, average (mm) | 0.75† | 0.85* | 0.001 | 0.73† | 0.82* | 0.015 |
| Vessel diameter (mm) | 5.85† | 6.4* | 0.006 | 5.90† | 6.20* | 0.134 |
| VASCUQOL-6 | 24† | 15† | <0.001 | 24† | 16† | <0.001 |
| Triglycerides (mmol/L) | 0.9* | 1.2† | 0.017 | 0.9* | 1.2† | 0.016 |
| Cholesterol (mmol/L) | 5.5† | 4‡ | <0.001 | 5.25† | 4† | <0.001 |
| HDL- Cholesterol (mmol/L) | 1.7† | 1.3‡ | <0.001 | 1.75† | 1.2† | <0.001 |
| LDL- Cholesterol (mmol/L) | 3.4† | 2.2§ | <0.001 | 3.3† | 2.35† | <0.001 |
| **Medical history (Y/N)** |  |  |  |  |  |  |
| Hypertension | 7/23 | 25/0 |  | 7/20 | 24/0 |  |
| Hyperlipidemia | 4/23 | 37/0 |  | 3/24 | 24/0 |  |
| Previous vascular surgery | 0/30 | 14/10 |  | 0/27 | 13/10 |  |
| Heart failure | 0/30 | 3/21 |  | 0/27 | 3/20 |  |
| Previous stroke | 0/30 | 5/19 |  | 0/27 | 5/18 |  |
| Ischemic heart disease | 0/30 | 7/17 |  | 0/27 | 7/16 |  |
| Hypertension medication | 5/24 | 21/2 |  | 5/20 | 20/2 |  |
| Antiarrhythmics and blood thinning medication | 3/27 | 21/3 |  | 3/23 | 19/3 |  |
| Statins | 3/27 | 20/3 |  | 2/24 | 18/3 |  |
| Other lipid-lowering drugs | 2/28 | 5/17 |  | 2/24 | 5/16 |  |
| Anti-inflammatory treatment/NSAIDs | 5/33 | 5/30 |  | 5/21 | 2/20 |  |
| Immunosuppressive therapy | 0/38 | 0/37 |  | 0/26 | 0/24 |  |
|  | *n=37 | *n=36 |  | * n=25 | * n= 23 |  |
|  | †n=38 | †n=34 |  | † n= 26 | † n=22 |  |
|  |  | ‡n=35 |  |  |  |  |
|  |  | §n=33 |  |  |  |  |

M/F: male female; ABI: ankle brachial index; VASCUQOL-6: Vascular Quality of Life Questionnaire-6; Y/N: yes/ no

During recruitment, the participants underwent various non-invasive examinations to assess cardiovascular risk by evaluating microvascular status, vessel diameter and intima media thickness (IMT). Ankle-brachial index (ABI) measurement with pen-Doppler system was performed to assess microvascular status in limbs. IMT of the vessels was evaluated by performing ultrasound of the carotid arteries at the physiology clinic, University Hospital Örebro. To assess the quality of life we have used a PAD-specific health related quality of life questionnaire VascuQoL6. This questionnaire is used in clinical practice for evaluation and in the national quality register SWEDVASC in Sweden.

The present study was carried out in accordance with the principles outlined in the Declaration of Helsinki and was approved by the Swedish Ethical Review Authority (Dnr 2021-03386). Written informed consent was obtained from all the participants.

### Preparation of blood samples

From each study participant, whole venous blood was drawn after a fasting period of at least 12 hours. Whole blood samples were either collected in sodium heparin, EDTA or lithium heparin tubes (BD Biosciences, Franklin Lakes, NJ, USA) to perform different analyses. Blood collected in sodium heparin and EDTA tubes were transported immediately at room temperature to our lab and the lithium heparin tubes to the clinical chemistry lab at University Hospital Örebro for traditional cardiovascular risk marker analysis.

### Soluble immune checkpoints

Soluble immune checkpoints were measured with the 14 plex ProcartaPlex™ Human Immuno-Oncology Checkpoint Panel 1 (Invitrogen, Waltham, MA, USA) using Bio-Plex^®^ 200 technology (Bio-RAD, Hercules, CA, USA), according to the manufacturer’s instructions. To this end, fasting blood plasma collected in EDTA tubes from 39 HCs and 37 PAD patients was used. The panel allows for the simultaneous detection of the co-stimulatory immune checkpoints CD27, CD28, CD80, CD137, GITR and herpesvirus entry mediator (HVEM). In addition, the concentration of co-inhibitory checkpoints BTLA, cytotoxic T-lymphocyte associated protein 4 (CTLA-4), indoleamine 2,3-dioxygenase (IDO), LAG-3, PD-1, PD-L1 and PD-L2, and TIM-3 can be assessed. Samples were analyzed in duplicate, and the levels of different checkpoint proteins were expressed in pg/mL.

### Surface staining membrane-bound immune checkpoints

Next, membrane-bound immune checkpoint expression was measured in sodium heparin blood of 27 HCs and 24 PAD patients by flow cytometry. First, we analyzed the expression of PD-L2 on monocyte subsets (classical: CD14+CD16-, intermediate: CD16dimCD14dim and non-classical: CD14-CD16+) and conventional dendritic cells (cDCs, CD16-CD14-CD11c+). Then, we analyzed the expression of CD28, BTLA, PD-1, TIM-3, LAG-3 and GITR on CD4+ and CD8+ T cells. The distribution of CD4+ and CD8+ differentiation subsets (naïve: CCR7+CD45RA+, central memory: CCR7+CD45RA-, effector memory: CCR7-CD45-, and terminally differentiated: CCR7-CD45RA+) was determined in a subset of patients (n=18) and controls (n=19). Flow cytometry staining was performed in blood incubated for 15 minutes at room temperature with monoclonal fluorescent antibodies (Supplementary table 1). Blood was subsequently incubated for 10 minutes with FACS Lysing solution (1X, BD Biosciences). After centrifugation, the samples were washed with PBS + 1% BSA and resuspended in 300 μL PBS + 1% BSA. Samples were analyzed on a Gallios Flow cytometer (Beckman Coulter, Brea, CA, USA). Fluorescence minus one (FMO) controls and unstained controls were used to set the gates (Gating strategy: Supplementary figure 1A-B).

### Cytokine production

To determine whether T cells of HCs and PAD patients that expressed markers for immune exhaustion were indeed exhausted, whole sodium heparin blood of 8 HCs and 13 PAD patients was stimulated to measure intracellular cytokine production with flow cytometry. Sodium heparin blood was diluted 1:1 with RPMI 1640 medium (Gibco, Waltham, Massachusetts, US) and stimulated with phorbol 12-myristate 13-acetate (PMA, 50 ng/mL, Sigma-Aldrich, St. Louis, MO, USA), ionomycin (1,6 μg/mL, Sigma-Aldrich) and brefeldin A (BFA, 1 μg/mL, BD Biosciences), for four hours at 37°C + 5% CO2. Control samples were only incubated with BFA. After four hours of stimulation, samples were mixed with Pharm lyse (1X, BD Biosciences) and incubated for 10 minutes at room temperature. Cells were washed once with FACS buffer (PBS + 2% fetal bovine serum (FBS), 2 mM EDTA) and then incubated with diluted fixable viability stain 510 (1:10, BD Biosciences) for 15 minutes. Samples were then stained for surface markers (CD3, CD4, CD8, PD-1 and TIM-3, Supplementary figure 1C, Supplementary table 1) for 15 minutes. Afterwards, samples were washed once with FACS buffer and fixed and permeabilized by adding 500 μL fixation/permeabilization solution (BD Biosciences). After 20 minutes of incubation, samples were centrifuged and then washed and incubated for 10 minutes with Perm/Wash buffer (1X, BD Biosciences). Samples were stained for intracellular cytokines (TNF-α, IL-17A, IL-10 and IFN-γ) for 30 minutes and washed again with Perm/Wash buffer. Samples were resuspended in FACS buffer and analyzed on a Gallios Flow cytometer (Beckman Coulter).

### Data analysis

Flow cytometry data was analyzed using Kaluza version 2.1 (Beckman coulter). The levels of soluble immune checkpoints were analyzed using Bio-Plex manager software version 6.2. Results were derived through comparison to a standard curve (5-parameter logistic curve fit) of known concentration of each analyte. Graphs were created and statistical analyses were executed in GraphPad Prism version 10. Statistical significance between HCs and PAD patients was tested using Mann-Whitney U tests. Statistical significance within the groups was determined with Wilcoxon signed-rank tests. For correlation analyses Spearman tests were used. P-values <0.05 were considered statistically significant.

## Results

### Patient population characteristics

We included 39 HCs and 37 PAD patients to assess the concentration of soluble immune checkpoints in blood plasma. The PAD patients were older in age than the HCs (p<0.001) and contained more active smokers than HCs. The microvascular status is significantly altered in PAD patients as the median ABI is lower in PAD patients than HCs (Table 1). The median vessel diameter of carotid arteries and maximum and average intima media thickness was higher in PAD patients than HCs (Table 1). PAD patients had a significantly lower VascuQoL6 score compared to HCs suggesting a declined quality of life in PAD patients (Table 1). All PAD patients experienced hypertension and hyperlipidemia, for which most of them are treated with hypertension medication and lipid lowering drugs. Only a small proportion of HCs experienced hypertension and hyperlipidemia. A large number of HCs (n=27) and PAD (n=24) were included in follow-up experiments in which we assessed the expression of membrane-bound immune checkpoints. In this subset of patients and controls, the age was not statistically significant different between the groups (p=0.057), as well as the vessel diameter. Other differences in clinical characteristics were similar as in the larger subset of patients and controls (Table 1).

### Soluble TIM-3 and PD-L2 concentrations differed between female PAD patients and HCs

We assessed the concentration of soluble immune checkpoints using Bioplex technology in blood plasma (Figure 1). The concentration of immune checkpoints was similar between patients and HCs (Figure 1A), although large variations could be observed within the groups. The concentration of PD-L1 was too low to be measured in blood plasma and is therefore not shown. Upon stratifying the groups based on sex, we found that female PAD patients had higher concentrations of soluble TIM-3 than female HCs (trend p=0,06), whereas the concentration of soluble TIM-3 was not different between male HCs and PAD patients (Figure 1B). Soluble PD-L2 was lower in female PAD patients than HCs and was similar between male HCs and PAD patients (Figure 1B).

**Figure 1:**
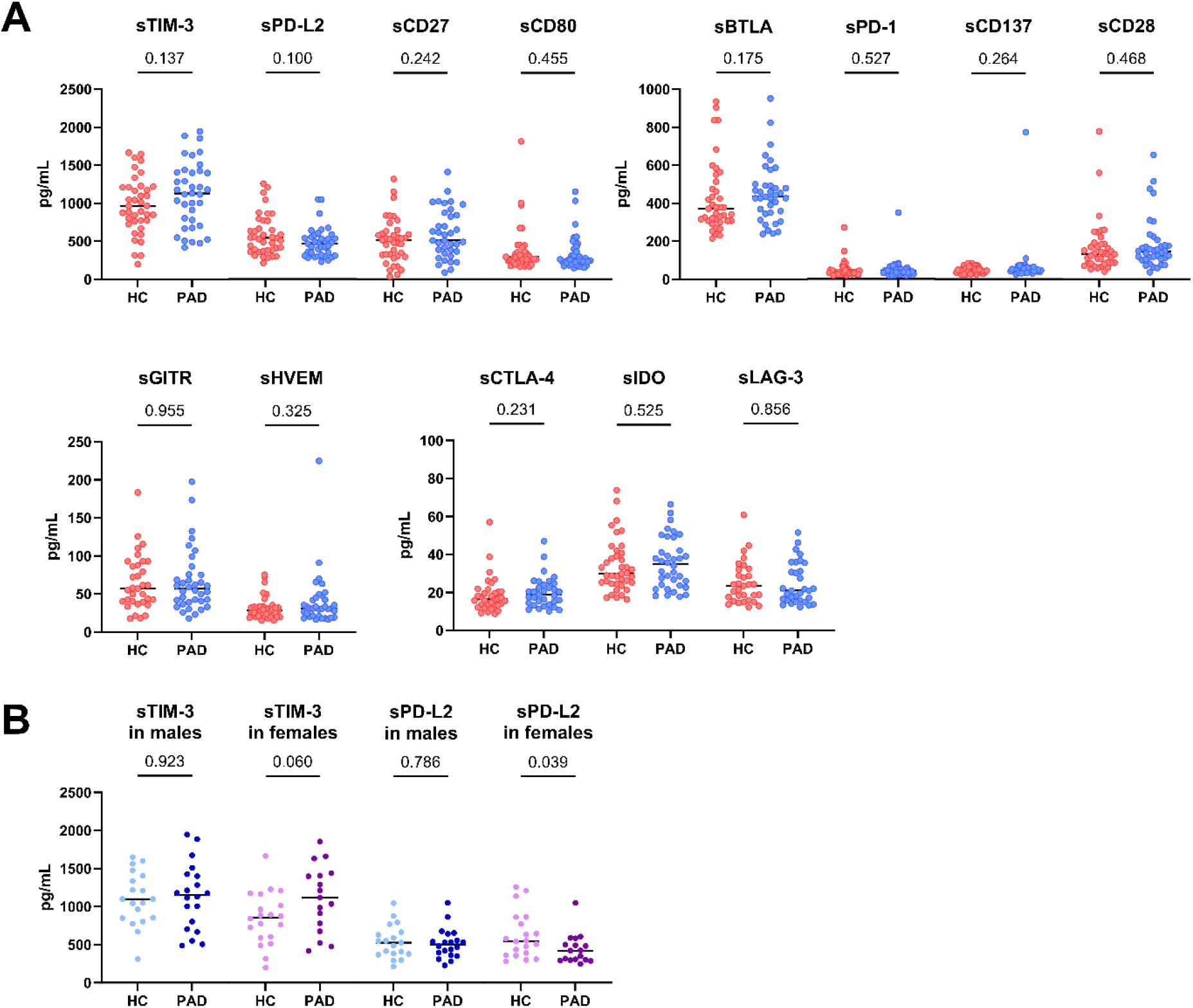
Concentration of soluble immune checkpoints in PAD and HCs. (A) The concentration of the measured soluble immune checkpoints was similar in PAD patients (n=37) and HCs (n=39). (B) The concentration of soluble TIM-3 was similar between male HCs (n=19) and PAD patients (n=20) but higher in female PAD patients (n=17) than HCs (n=20). The concentration of soluble PD-L2 was similar between males but lower in female PAD patients than HCs. Horizontal bars reflect the median. Mann-Whitney U tests were used to compare HCs and PAD patients. P-values are shown in the graphs. PAD: peripheral artery disease. HC: healthy controls. TIM-3: T-cell immunoglobulin and mucin domain 3. PD-L2: programmed cell death 1 ligand 2. BTLA: B-and T-lymphocyte attenuator. PD-1: programmed cell death 1. GITR: glucocorticoid-induced TNRF-related protein. HVEM: herpesvirus entry mediator. CTLA-4: cytotoxic T-lymphocyte associated protein 4. IDO: indoleamine 2,3-dioxygenase. LAG-3: lymphocyte-activation gene 3.

Since the PAD group was significantly older than the HCs group, we assessed whether age correlated with soluble PD-L2 and TIM-3 concentrations. We also assessed whether these checkpoints correlated with clinical characteristics. Age did not correlate with either soluble TIM-3 or PD-L2 concentrations in PAD patients. Correlation analyses of soluble PD-L2 concentration and clinical characteristics showed that soluble PD-L2 positively correlated with average intima media thickness (R=0,34, p=0,046). Soluble TIM-3 concentrations did not correlate with clinical characteristics.

### Increased proportions of membrane-bound PD-L2 on monocytes, TIM-3 on CD4+ and LAG-3 on CD8+ T cells in PAD patients

As soluble PD-L2 and TIM-3 concentrations were different between female PAD patients and HCs, we assessed the expression of their membrane-bound forms in APCs and T cells. Furthermore, we assessed the expression of PD-1, GITR, LAG-3 and BTLA on T cells as well, as these have previously been shown to have implications in atherosclerosis, as mentioned earlier. CD28 was included as well as lack of CD28 indicates immune ageing/ senescence.

The distribution of monocyte-subsets was different between PAD patients and HCs (Figure 2A). The frequencies of classical monocytes were higher in PAD patients than in HCs. Intermediate monocyte-frequencies were similar between the groups and the frequency of non-classical monocytes was lower in PAD patients. Within each monocyte subset, the frequencies of cells expressing PD-L2 were higher in PAD patients than in HCs (Figure 2B). This pattern was similar in males and females between HCs and PAD patients, but the analyses lacked statistical power due to the smaller group sizes (Supplementary figure 2). PD-L2+ frequencies within conventional DCs were similar between PAD and HCs (Supplementary figure 3).

**Figure 2:**
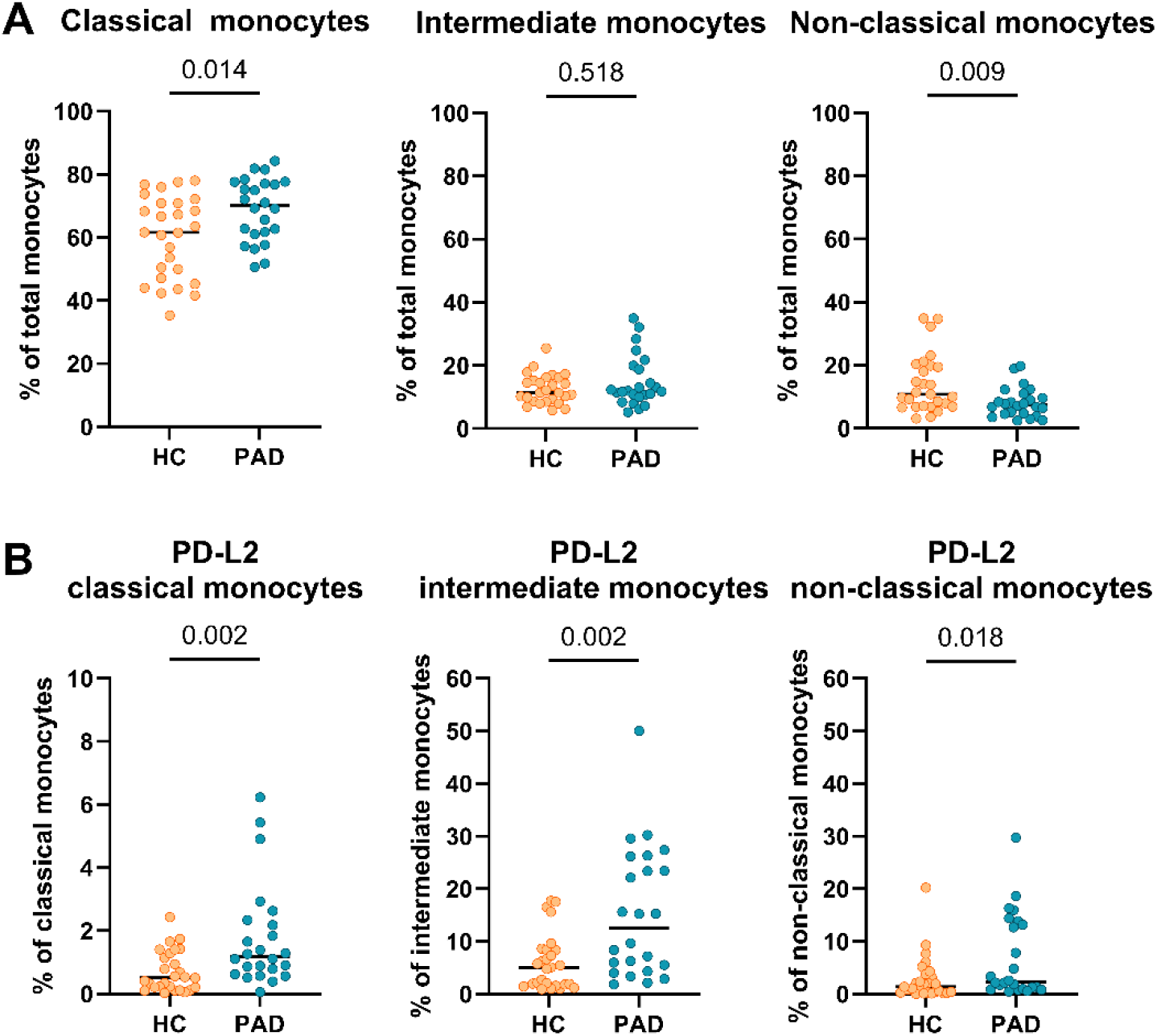
Frequencies of monocytes and expression of PD-L2 within monocytes subsets. (A) The frequency of classical monocytes was higher, and the frequency of non-classical monocytes was lower in PAD patients (n=24) than HCs (n=27). (B) PD-L2 frequencies were higher within all monocyte subsets in PAD patients than in HCs. Horizontal bars reflect the median. Mann-Whitney U tests were used to compare HCs and PAD patients. P-values are shown in the graphs. PAD: peripheral artery disease. HC: healthy controls. PD-L2: programmed cell death 1 ligand 2.

Next, we assessed the expression of immune checkpoint receptors on CD8+ and CD4+ T cells. We found the frequencies of cells expressing CD28, BTLA, PD-1 and GITR to be similar within CD8+ and CD4+ T cells of HCs and PAD patients (Figure 3). The frequency of TIM-3-expressing cells within CD8+ T cells was similar between HCs and PAD patients but higher in CD4+ T cells of PAD patients (Figure 3). Although the difference in soluble TIM-3 concentration was mainly found between female HCs and PAD patients, we did not find such as trend regarding membrane-bound TIM-3 expression. Finally, LAG-3 was virtually absent on CD8+ T cells of HCs, and found to be higher among CD8+ T cells of PAD patients (Figure 3). LAG-3 also correlated negatively with vessel diameter in PAD patients (R=-0,41, P=0,05).

**Figure 3:**
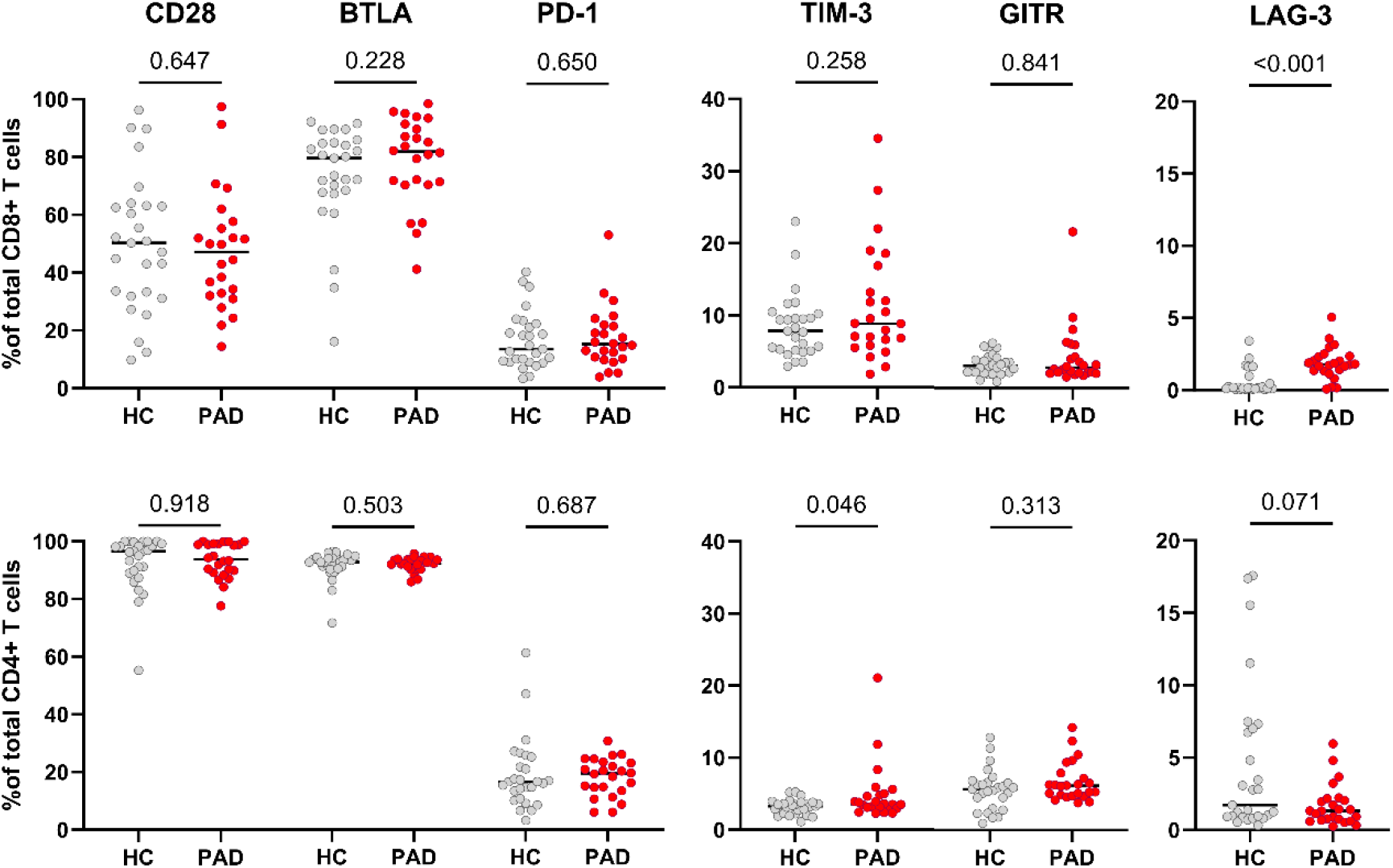
Expression of membrane-bound immune checkpoint receptors in CD8+ and CD4+ T cells of HCs and PAD patients. The upper panel shows the frequencies of cells expressing CD28, BTLA, PD-1, TIM-3, GITR and LAG-3 within CD8+ T cells of HCs (n=27) and PAD patients (n=24). The lower panel shows the frequencies of these checkpoints within CD4+ T cells. Horizontal bars reflect the median. Mann-Whitney U tests were used to compare HCs and PAD patients. P-values are shown in the graphs. PAD: peripheral artery disease. HC: healthy controls. BTLA: B-and T-lymphocyte attenuator. PD-1: programmed cell death 1. TIM-3: T-cell immunoglobulin and mucin domain 3. GITR: glucocorticoid-induced TNRF-related protein. LAG-3: lymphocyte-activation gene 3.

As the expression of immune checkpoints can differ within CD4+ and CD8+ T cell differentiation subsets, we analyzed in a smaller randomly selected proportion of patients and controls whether the distribution of differentiation subsets was changed. We observed no differences in either CD4+ or CD8+ differentiation subsets (Supplementary table 2).

### Co-expression analyses of immune checkpoints reveal differences between HCs and PAD patients

As the frequency of TIM-3+ cells within CD4+ T cells was higher in PAD patients than in HCs, we assessed whether co-expression with other checkpoints was different between the groups as well. We found that the majority of TIM-3+ CD4+ T cells were negative for LAG-3, and this was even more pronounced in PAD patients. The relative frequency of TIM3+GITR+ was lower in PAD patients than in HCs. Around 20% of TIM-3+ cells expressed PD-1 as well, and almost all TIM-3+ cells were also positive for CD28 and BTLA (figure 4).

**Figure 4:**
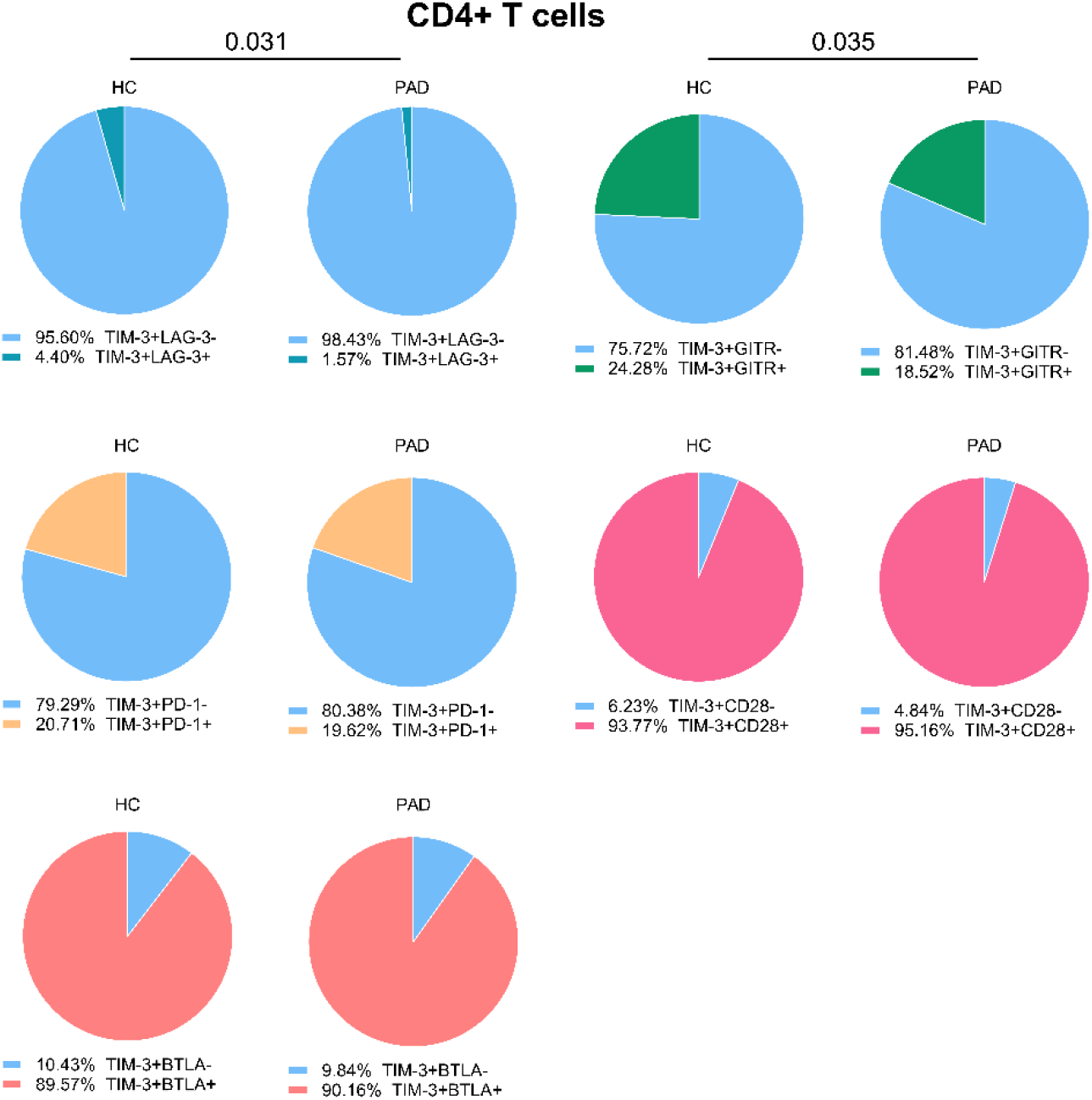
Co-expression of TIM-3 with other immune checkpoints in CD4+ T cells. We show the relative co-expression of TIM-3+ cells with other immune checkpoints within CD4+ T cells in PAD patients (n=24) and HCs (n=27). Shown are the frequencies of total TIM-3+ cells that do or do not express the other indicated immune checkpoints. Mann-Whitney U tests were used to compare HCs and PAD patients. P-values are shown in the graphs. PAD: peripheral artery disease. HC: healthy controls. BTLA: B-and T-lymphocyte attenuator. PD-1: programmed cell death 1. TIM-3: T-cell immunoglobulin and mucin domain 3. GITR: glucocorticoid-induced TNRF-related protein. LAG-3: lymphocyte-activation gene 3.

Previous analyses show that CD8+ T cells have a higher frequency of LAG-3 expressing cells in PAD patients than in HCs. The large majority of LAG-3+ cells were TIM-3 negative, although the relative percentage of LAG-3+TIM-3+ CD8+ T cells was slightly higher in PAD patients. LAG-3+ CD8+ T cells were largely negative for GITR and PD-1 expression as well. Around half of the LAG-3+ CD8+ T cells co-expressed CD28 and almost all LAG-3+ cells expressed BTLA (Supplementary figure 4). These results indicate that exhaustion/inhibition markers such as PD-1, TIM-3, BTLA and LAG-3 are not all co-expressed on the same cells.

### TIM-3+ CD4+ T cells are exhausted in PAD patients and HCs

Our results show that the frequency of TIM-3+CD4+ T cells was higher in PAD patients, and that a small frequency co-expressed exhaustion markers such as LAG-3 and PD-1, but a majority expressed BTLA and CD28. We therefore analyzed whether TIM-3+ cells are exhausted, as defined by the reduced capacity to produce (pro)-inflammatory cytokines. We show that TIM-3+CD4+ T cells of PAD patients and HCs had similar cytokine production capacity (Figure 5A). Compared to TIM-3-CD4+ T cells, TIM-3+CD4+ T cells of both PAD patients and HCs had lower frequencies of cells expressing TNF-α, IFN-γ and IL-17A. However, they had a relatively higher frequency of IL-10+ cells (Figure 5B). This indicates that a large subset of TIM-3+ CD4+ T cells is indeed exhausted.

**Figure 5.**
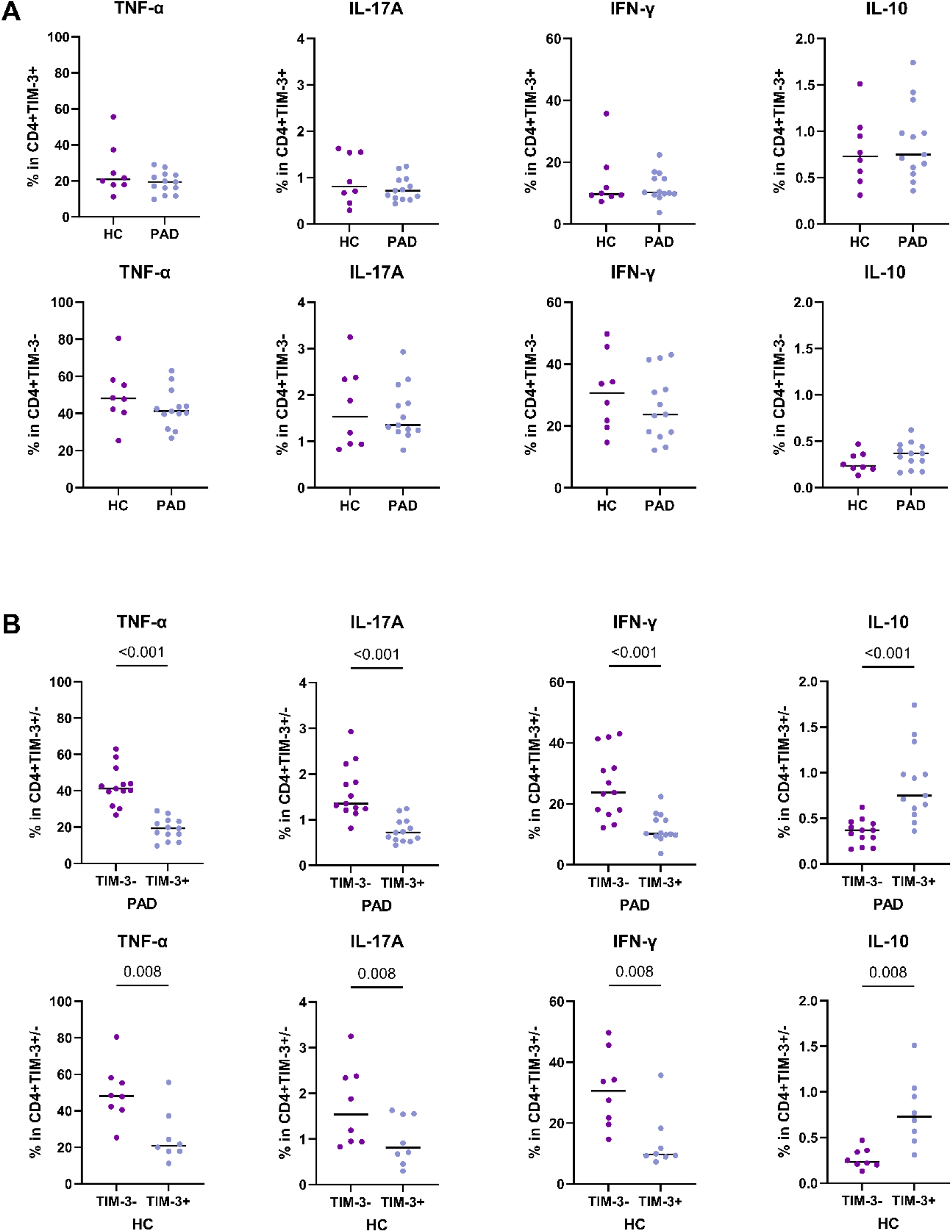
Cytokine production by TIM-3+ CD4+ T cells. Intracellular cytokine production after stimulation of CD4+ T cells was assessed in PAD patients (n=13) and HCs (n=8). (A) The total frequency of TIM-3+CD4+ T cells expressing intracellular TNF-α, IFN-γ, IL-17A and IL-10 are shown in the upper panel. The lower panel shows the frequency of TIM-3-CD4+ T cells expressing intracellular TNF-α, IFN-γ, IL-17A and IL-10. B) The pair-wise comparison of intracellular TNF-α, IFN-γ, IL-17A and IL-10-expressing cells within TIM-3- and TIM-3+ CD4+ T cells is shown within the PAD patients and HCs. Horizontal bars reflect the median. Mann-Whitney U tests were used to compare HCs and PAD patients. Wilcoxon signed-rank tests were used to compare within the groups. P-values are shown in the graphs. PAD: peripheral artery disease. HC: healthy controls. TIM-3: T-cell immunoglobulin and mucin domain 3. TNF-α: Tumor necrosis factor alpha, IFN-γ: Interferon gamma.

## Discussion

In this study, we aimed to investigate whether concentrations of soluble immune checkpoints and the expression of membrane-bound immune checkpoints were different between PAD patients and HCs. We found that soluble immune checkpoints and membrane-bound immune checkpoints show different expression patterns between PAD patients and HCs.

At first glance, soluble immune checkpoints concentrations appear to be largely similar between PAD patients and HCs. However, after further analysis we show that there is a sex-difference in soluble immune checkpoint concentrations between PAD patients and HCs, as soluble PD-L2 was lower in female PAD patients and soluble TIM-3 showed a trend to be higher in female PAD patients than female HCs. The effects of sex on immune checkpoint expression have been shown before in young and older healthy donors, especially regarding PD-1 expression (15). We are not aware of studies showing differences between females and males regarding soluble immune checkpoint concentrations. Although soluble levels of PD-L2 were lower in female PAD patients, soluble PD-L2 positively correlated with intima media thickness. Intima media thickness is a marker for disease severity, but since the levels of PD-L2 were lower in female PAD patients, it is challenging to interpret this correlation.

Next, we show that the frequency of membrane-bound PD-L2-expressing cells was higher in monocyte subsets of PAD patients. Membrane-bound PD-L2 expression frequencies were not different between female and male patients and controls. In summary, the results suggest that there are differences in the post-transcriptional processing of PD-L2 that lead to increased surface expression but reduced plasma concentration in PAD.

The role of the PD-1-PD-L1/PD-L2 axis in atherosclerosis is still up to debate. A previous study showed that the PD-1-PD-L1/PD-L2 axis was protective in preventing atherosclerotic lesion formation in mice (16). However, another study in human adult cancer patients with atherosclerotic plaques showed that targeting PD-1 reduces atherosclerotic plaque size. Plaque-specific PD-1+ T cells appeared to be pro-inflammatory instead of exhausted. These cells also did not express other markers of exhaustion, such as LAG-3 and TIM-3. The authors stated that this pro-inflammatory state could be caused by the lack of PD-L1 and PD-L2 expression in the atherosclerotic plaque. Without being able to bind to PD-1 ligands, PD-1+ T cells will not receive inhibitory signals that will lead to immune suppression. The monoclonal antibody acted as a substitute PD-1 ligand, which suppressed the pro-inflammatory functions of PD-1+ T cells, thereby reducing plaque size (17). This study emphasizes that it would be important to investigate PD-1 and PD-L1/PD-L2 expression in plaques from PAD patients as well, to assess whether the local tissue expression of these immune checkpoints differ from our findings in blood.

Further inspection of immune checkpoint expression on T cells revealed that the frequency of TIM-3+ was higher in CD4+ T cells whereas the frequency of LAG-3+ cells was higher in CD8+ T cells of PAD patients. LAG-3 has previously been targeted in an atherosclerotic mouse model. Here, LAG-3 knockout mice or LAG-3 blockade with monoclonal antibodies had no effects on plaque size. However, the levels of pro-inflammatory CD4+ T cells increased and accumulated more in plaques of mice lacking LAG-3 (18). LAG-3 could therefore be an important regulator of T cell activation in atherosclerosis. In our study, the increased expression could indicate a regulatory mechanism in PAD patients to counteract possible increased CD8+ T cell responses. This is also emphasized by the fact that LAG-3 correlated negatively with vessel diameter in PAD patients.

TIM-3 expression was increased in CD4+ T cells of PAD patients. In a mouse model of atherosclerosis, blocking TIM-3 reduced lesion size and increased monocyte and CD4+ T cell frequencies (19). In PAD patients with intermittent claudication, only an increase in PD-1+TIM-3+CD8+ T cells was reported (12). This study was performed in patients before start of treatment, which could potentially explain the differences in outcome with our study. Another study showed that expression of TIM-3+CD4+ T cells was increased in coronary heart disease patients as well and correlated with disease severity (20). In our study, the expression of TIM-3 did not correlate with clinical characteristics of PAD patients. We also show that a large percentage of TIM-3+CD4+ T cells lacked expression of other exhaustion markers, both in PAD patients as in HCs. However, a large percentage of TIM-3+ cells showed signs of exhaustion. The increased expression in PAD could therefore be a consequence of prolonged immune activation.

The main limitation of our study is that we included PAD patients that are currently being treated for hypertension and hyperlipidemia. Therefore, we could not determine whether immune checkpoint expression was associated with markers of hypertension or lipid levels. Furthermore, we have not included plaque-specific immune cells. In future studies, the expression of immune checkpoints in plaques of PAD patients should be determined, to learn more about local pathogenesis in PAD.

In conclusion, we show that PAD patients show differences in immune checkpoint expression of both membrane-bound and soluble immune checkpoints, especially related to markers of immune exhaustion. Although exhaustion markers did not always overlap, the increase in exhaustion markers might be caused by prolonged immune activation. Future investigations should be focused on immune checkpoint expression in atherosclerotic lesions in PAD patients. Furthermore, it would be interesting to investigate whether immune cells of PAD patients with critical limb ischemia show even more signs of immune exhaustion.

## Acknowledgements

We are very grateful to the staff, especially Christine Degner at the Clinical Trial Unit, University Hospital Örebro, for their excellent support in collecting the blood samples. We also thank the biomedical analysts, Frida Tornqvist and Gabriella Persson, from the physiology clinic, University Hospital Örebro, for their assistance in the physiological assessments in this study.

## Author Contribution

RR, SK, AKK conceptualized the study. Experiments were conducted by RR and SK. Data was analyzed by RR and SK. The original draft was written by RR and AKK. MD contributed to patient recruitment and clinical information. IR, HH, MD and AS contributed to the interpretation of the results. All authors were involved in reviewing and editing the manuscript and approved the final draft.

