## Supplementary tables and figures for "Patients with peripheral artery disease demonstrate altered expression of soluble and membrane-bound immune checkpoints by peripheral blood immune cells"

Supplementary Table 1: Overview of the antibodies that were used in this study

| Marker | Fluorochrome | Clone | Catalogue number | Company |
|---|---|---|---|---|
| CD19 | R718 | HIB19 | 567343 | BD Biosciences |
| CD56 | R718 | B159 | 566965 | BD Biosciences |
| CD3 | R718 | OKT3 | 567348 | BD Biosciences |
| CD15 | AF700 | SSEA-1 | 301920 | Biolegend |
| HLA-DR | BV510 | G46-6 | 563083 | BD Biosciences |
| CD14 | APC-Cy7 | MϕP-9 | 557831 | BD Biosciences |
| CD16 | PE-CF594 | 3G8 | 562293 | BD Biosciences |
| CD11c | PerCP-Cy5.5 | B-ly6 | 565227 | BD Biosciences |
| PD-L2 | PE | MIH18 | 558066 | BD Biosciences |
| CD3 | APC-H7 | SK7 | 560176 | BD Biosciences |
| CD4 | PE-CF594 | RPA-T4 | 562281 | BD Biosciences |
| CD8 | PE-Cy7 | RPA-T8 | 557746 | BD Biosciences |
| GITR | BV510 | V27-580 | 747665 | BD Biosciences |
| Lag-3 | PerCP-Cy5.5 | 11C3C65 | 369312 | Biolegend |
| CD28 | BB515 | CD28.2 | 564492 | BD Biosciences |
| PD-1 | APC | MIH4 | 558694 | BD Biosciences |
| TIM-3 | BV421 | 7D3 | 565562 | BD Biosciences |
| BTLA | PE | J168-540 | 558485 | BD Biosciences |
| TNF-α | PerCP-Cy5.5 | MAb11 | 560679 | BD Biosciences |
| IL-17A | Alexa Fluor 488 | N49-653 | 560488 | BD Biosciences |
| IL-10 | PE | JES3-19F1 | 559330 | BD Biosciences |
| IFN-γ | R718 | 4S.B3 | 567060 | BD Biosciences |
| CD45RA | FITC | HI100 | 555488 | BD Biosciences |
| CD196(CCR6) | PE | QA17A37 | 399004 | Biolegend |
| CD197 (CCR7) | PE-CF594 | 2-L1-A | 566768 | BD Biosciences |
| CD3 | PerCP-Cy5.5 | UCHT1 | 560835 | BD Biosciences |
| CD183 (CXCR3) | PE- Cy7 | G025H7 | 353720 | Biolegend |
| CD25 | APC | 2A3 | 340907 | BD Biosciences |
| CD8 | APC-R700 | RPA-T8 | 565165 | BD Biosciences |
| CD45 | APC- H7 | 2D1 | 560178 | BD Biosciences |
| CD4 | V450 | RPA-T4 | 560345 | BD Biosciences |
| Fixable viability stain (FVS) | 510 | | 564406 | BD Biosciences |

Supplementary Table 2: Distribution of memory and naïve CD4+ and CD8+ T cell subsets

| | HC n=19* | PAD n=18* |
|---|---|---|
| CD4+ T naive | 43.83 | 39.04 |
| CD4+ T central memory | 36.63 | 43.23 |
| CD4+ T effector memory | 14.27 | 10.26 |
| CD4+ T terminally differentiated | 1.56 | 1.1 |
| CD8+ T naive | 15.04 | 16.47 |
| CD8+ T central memory | 8.8 | 11.21 |
| CD8+ T effector memory | 17.55 | 14.45 |
| CD8+ T terminally differentiated | 55.49 | 57.94 |

**Median frequencies within total CD4+ or CD8+ T cells are shown*

**A**

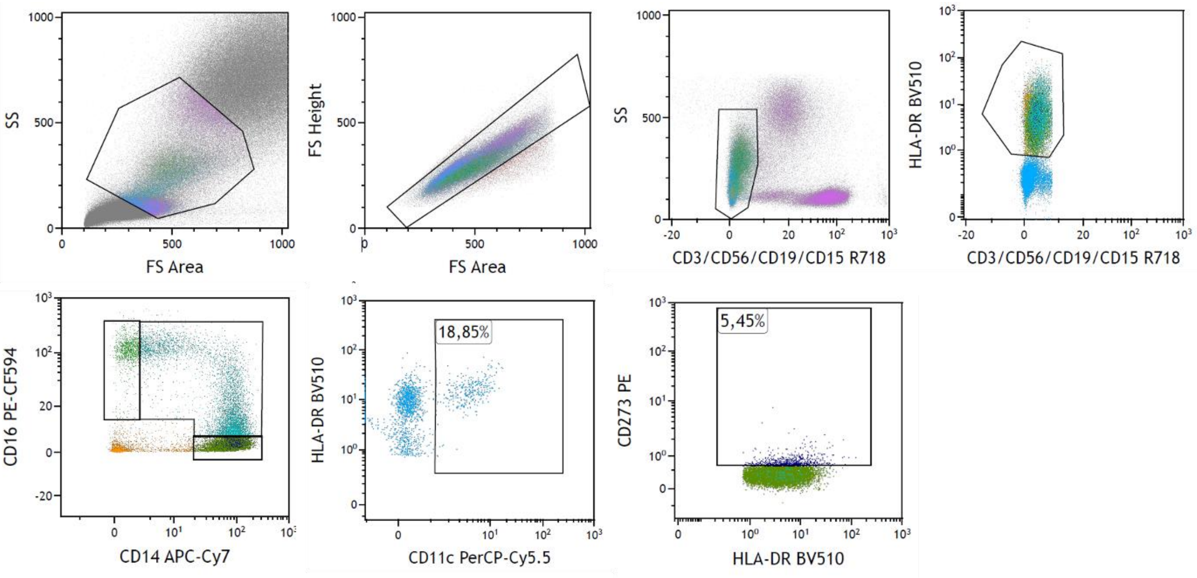
SS
FS Area
FS Height
CD3/CD56/CD19/CD15 R718
HLA-DR BV510
CD16 PE-CF594
CD14 APC-Cy7
CD11c PerCP-Cy5.5
18,85%
CD273 PE
5,45%

**B**

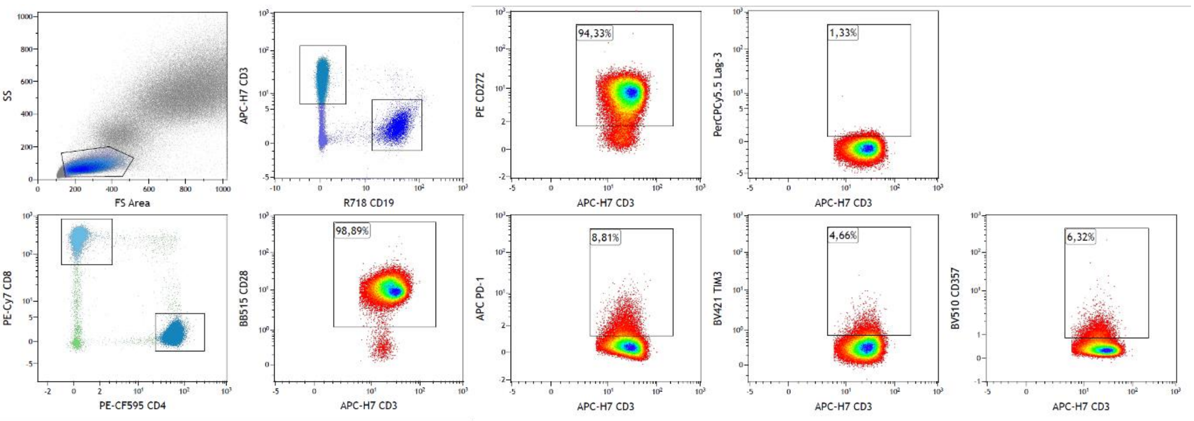
SS
FS Area
APC-H7 CD3
R718 CD19
PE CD272
94,33%
PerCPCy5.5 Lag-3
1,33%
PE-Cy7 CD8
PE-CF595 CD4
BB515 CD28
98,89%
APC PD-1
8,81%
BV421 TIM3
4,66%
BV510 CD357
6,32%

**C**

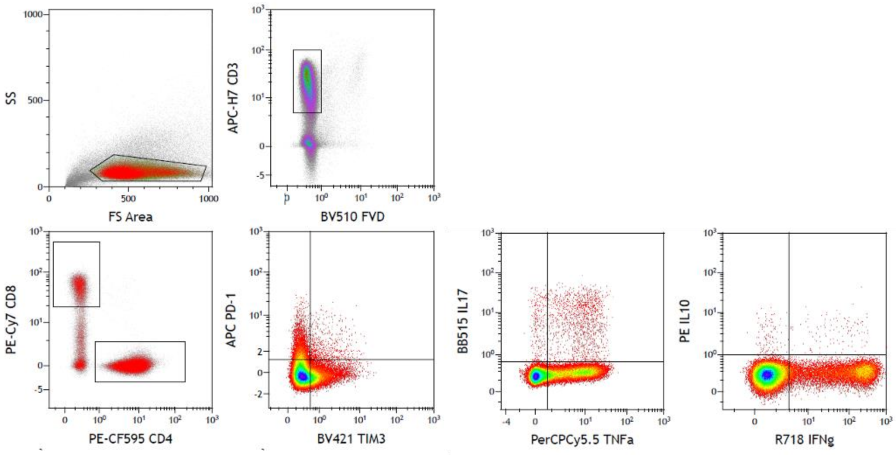
SS
FS Area
APC-H7 CD3
BV510 FVD
PE-Cy7 CD8
PE-CF595 CD4
APC PD-1
BV421 TIM3
BB515 IL17
PerCPCy5.5 TNFa
PE IL10
R718 IFNg

**Supplementary figure 1: Gating strategy.** (A) Membrane-bound immune checkpoint staining on antigen presenting cells. The forward and side scatter were used to select monocytes based on size and granularity. Single cells negative for CD3, CD56, CD19 and CD15 expression were further gated based on HLA-DR expression. Monocyte subsets were gated based on CD14 and CD16 expression into classical (CD14+CD16-), intermediate (CD16dimCD14dim) and non-classical (CD14-CD16+) monocytes. Conventional dendritic cells were CD16-CD14-CD11c+. PD-L2 expression was measured in each subset, and gating was based on the unstained. (B) Membrane-bound immune checkpoint staining on T cells. Lymphocytes were gated based on forward and side scatter, after which T cells were selected as CD3+ CD19-. Immune checkpoints were measured within CD4+CD3+ T cells and CD8+CD3+ T cells. Fluorescence minus one control (FMO) were used to set the gates and to check the overlap between colours. Gate settings for CD28, BTLA (CD272), LAG-3 and PD-1 were based on the expression of these checkpoints on B cells. GITR (CD357) gates were based on unstained cells and FMOs were used to set the gates for TIM-3+ cells. (C) Cytokine staining *in* vitro experiments. Gate settings for intracellular cytokine production analyses. Gates for the PD-1 and TIM-3 expression plots were based on B cells, to clearly distinguish between positive and negative populations. The gates for cytokine producing cells were based on the unstimulated samples. PAD: peripheral artery disease. HC: healthy controls. PD-L2: programmed cell death 1 ligand 2. BTLA: B-and T-lymphocyte attenuator. PD-1: programmed cell death 1. TIM-3: T-cell immunoglobulin and mucin domain 3. GITR: glucocorticoid-induced TNRF-related protein. LAG-3: lymphocyte-activation gene 3.

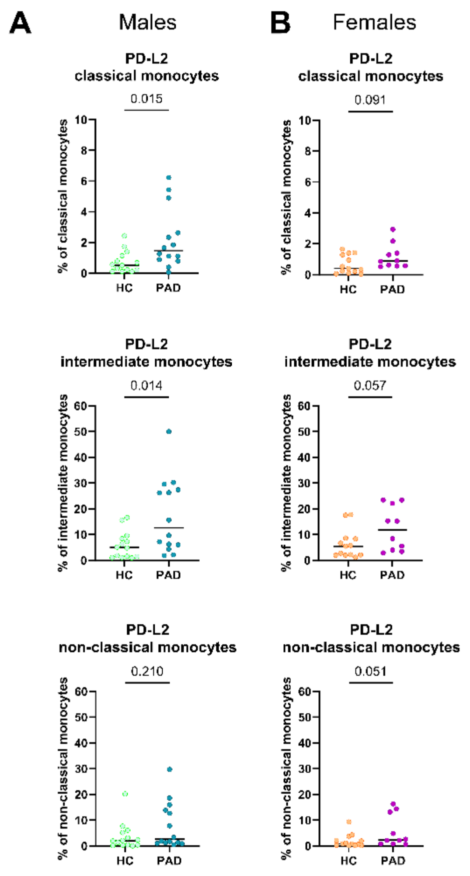


**Supplementary figure 2: Frequencies of PD-L2-expressing monocytes in female and male PAD patients and controls**. (A) Frequencies of PD-L2-expressing cells within classical monocytes, intermediate monocytes and non-classical monocytes in male HCs (n=14) and PAD patients (n=14). (B) Frequencies of PD-L2-expressing classical monocytes, intermediate monocytes and non-classical monocytes in female HCs (n=13) and PAD patients (n=10). Horizontal bars reflect the median. Mann-Whitney U tests were used to compare between HCs and PAD patients- P-values are shown in the graphs. PAD: peripheral artery disease. HC: healthy controls. PD-L2: programmed cell death 1 ligand 2.

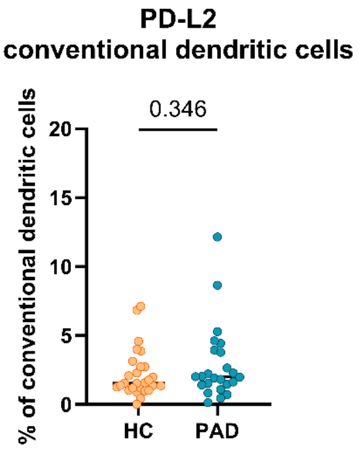


**Supplementary figure 3: PD-L2 expression on conventional dendritic cells.** Frequencies of conventional dendritic cells expressing PD-L2. Horizontal bars reflect the median. Mann-Whitney U tests were used to compare HCs (n=27) and PAD patients (n=24). P-values are shown in the graphs. PAD: peripheral artery disease. HC: healthy controls. PD-L2: programmed cell death 1 ligand 2.

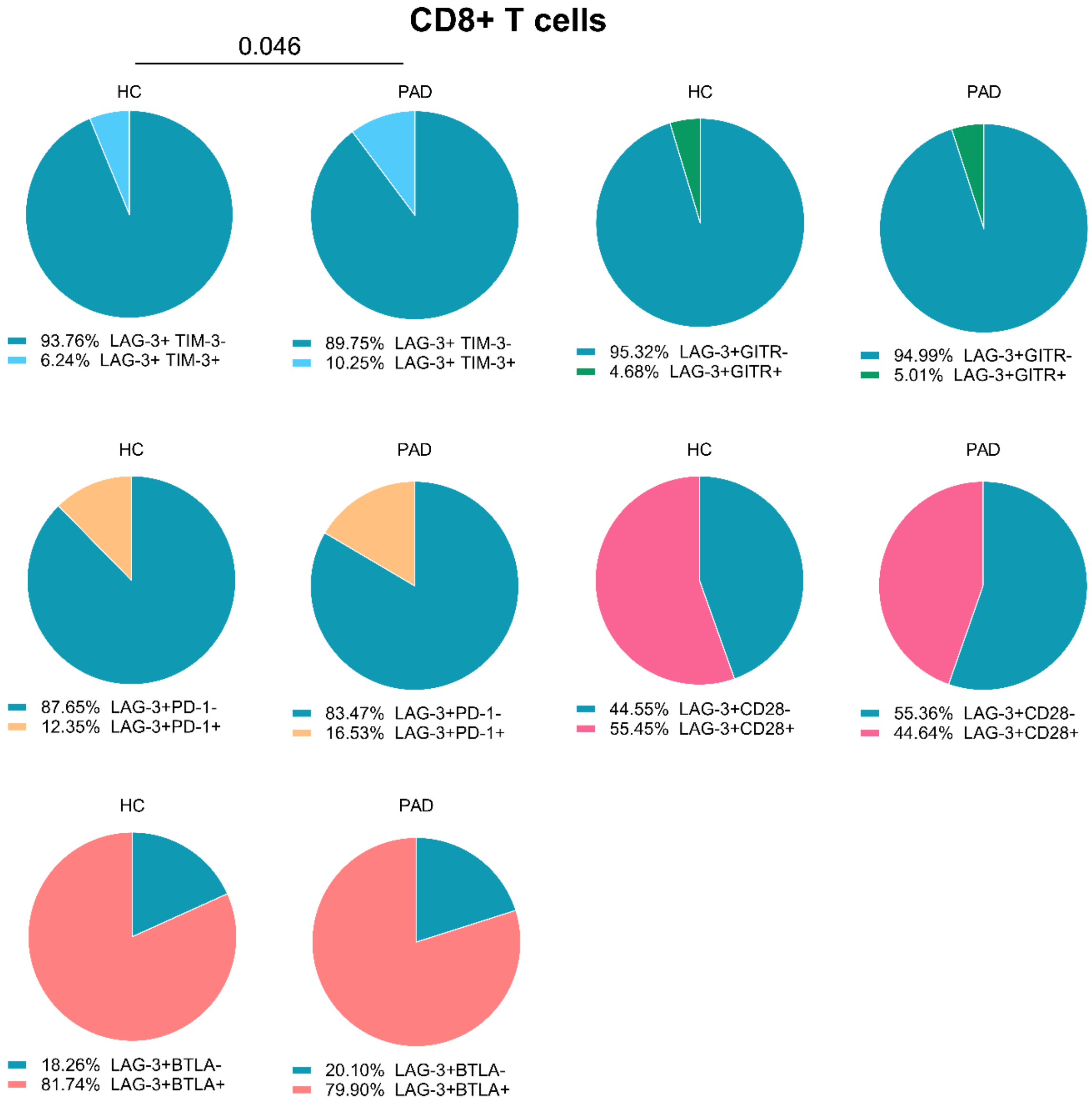


**Supplementary figure 4: Co-expression of LAG-3 with other immune checkpoints in CD8+ T cells.** We show the relative co-expression of LAG-3+ cells with other immune checkpoints within CD8+ T cells in PAD patients (n=24) and HCs (n=27). Shown are the frequencies of total LAG-3+ cells that do or do not express the other indicated immune checkpoints. Mann-Whitney U tests were used to compare HCs and PAD patients. P-values are shown in the graphs. PAD: peripheral artery disease. HC: healthy controls. BTLA: B-and T-lymphocyte attenuator. PD-1: programmed cell death 1. TIM-3: T-cell immunoglobulin and mucin domain 3. GITR: glucocorticoid-induced TNRF-related protein. LAG-3: lymphocyte-activation gene 3.
